# The homeodomain transcription factor Ventx2 regulates respiratory progenitor cell number and differentiation timing during *Xenopus* lung development

**DOI:** 10.1101/2022.06.13.495914

**Authors:** Scott A. Rankin, Aaron M. Zorn

## Abstract

Ventx2 is an *antennapedia* superfamily / NKL subclass homeodomain transcription factor best known for its role in the regulation of early dorsal-ventral pattern during Xenopus gastrulation and in the maintenance of neural crest multipotency. In this work we characterize an unappreciated spatial-temporal expression domain of *ventx2* in Xenopus respiratory system epithelial progenitors. We find *ventx2* is directly induced by BMP signaling in the ventral foregut prior to *nkx2-1*, the earliest epithelial marker of the respiratory lineage. Functional studies demonstrate that Ventx2 regulates the number of Nkx2-1/Sox9+ respiratory progenitors induced during foregut development, the timing and level of surfactant protein gene expression, and proper tracheal-esophageal separation. Our data suggest that Ventx2 regulates the balance of respiratory progenitor expansion and differentiation. While the *ventx* gene family has been lost from the mouse genome during evolution, humans have retained a *ventx2*-like gene *(VENTX)* and we lastly discuss how our findings might suggest a possible function of *VENTX* in human respiratory progenitors.

## Introduction

Ventx2 is an *antennapedia* superfamily / NK-like (NKL) subclass homeodomain (HD) transcription factor best known in Xenopus for its role in regulating early dorsal-ventral pattern during gastrulation (Sander et al 2007; McLin et al 2007; Rankin et al 2011). The NKL subclass of HD transcription factors (TFs) are named after Kim and Nirenberg who identified several when searching for homeobox genes in *Drosophila* (Kim and Nirenberg 1989). In Xenopus, the *ventx* family has 6 members (reviewed in Kumar et al 2022); the genes *ventx2*.*1* and *ventx2*.*2* have been extensively studied during blastula and gastrula stages where they are co-expressed as part of the BMP syn-expression group (Karaulanov et al 2004; von Bubnoff et al 2005; Paulsen et al 2011) and are also regulated by Wnt/B-catenin signaling (Hikasa et al 2010). Ventx2 proteins generally act as transcriptional repressors, functioning to promote ventral mesendoderm fates and antagonize the Spemann Organizer dorsal-anterior gene regulatory network in the early Xenopus embryo (Sander et al 2007; Rankin et al 2011; Rankin and Zorn 2022).

Recent work has also shown Ventx2 plays a key role in maintaining the multipotency of embryonic progenitor cells in the early blastula embryo (Scerbo et al 2012; Scerbo et al 2017) as well as maintaining the multipotency of developing neural crest cells during subsequent development (Scerbo et al 2020; Buitrago-Delgado et al 2015). The early function of Ventx2 in the blastula embryo has been suggested to be equivalent to the role mammalian Nanog plays in maintaining pluripotency of human and mouse embryonic stem cells (Scerbo et al 2012; reviewed in Kumar et al 2022).

The expression and function of Ventx2 later in Xenopus development, outside of the neural crest, has not been extensively studied. In this work, we characterize an unappreciated spatial-temporal expression domain of *ventx2* in Xenopus respiratory system epithelial progenitors. We find *ventx2* is directly induced by BMP signaling in the ventral foregut prior to *nkx2-1*, the earliest epithelial marker of the respiratory lineage. Functional studies demonstrate that Ventx2 regulates the number of Nkx2-1/Sox9+ respiratory progenitors induced during foregut development, the timing and level of surfactant protein gene expression, and proper tracheal-esophageal separation. Our data suggest that Ventx2 regulates the balance of respiratory progenitor expansion and differentiation. While the *ventx* gene family has been lost from the mouse genome during evolution, humans have retained a *ventx2*-like gene *(VENTX)* and we discuss the implications of our findings in regards to the possible function of *VENXT* in human respiratory progenitor cells.

## Results

### *ventx2* is expressed in respiratory epithelial progenitor cells prior to *nkx2-1*

In a search of gene expression patterns in the online biomedical knowledgebase Xenbase (*Xenbase*.*org*, Nenni et al. 2019) for genes that might serve as additional patterning markers during early Xenopus foregut and respiratory system development, we observed an unappreciated expression domain of the homeodomain TF *ventx2*.*1* in what appeared to be respiratory progenitors (images deposited by the Richard Harland lab: Xenbase gene page XB-GENEPAGE-919663, *Xenbase*.*org*). The genes *ventx2*.*1* and *ventx2*.*2* are highly similar, approximately 92% identical (Supplemental Fig.S1). Thus to characterize *ventx2* expression in detail, we performed in-situ hybridization on developing *X*.*tropicalis* embryos during endoderm organogenesis, encompassing gastrula through lung-bud stages, NF10.5 to NF43 (Fig.1, Supplemental Fig.S2), using a full length *ventx2*.*1* cDNA clone, which due to the high level of sequence identity should detect both *ventx2*.*1* and *ventx2*.*2*. In this work we thus refer to *ventx2*.*1/ventx2*.*2* collectively as *ventx2*.

**Figure 1.**
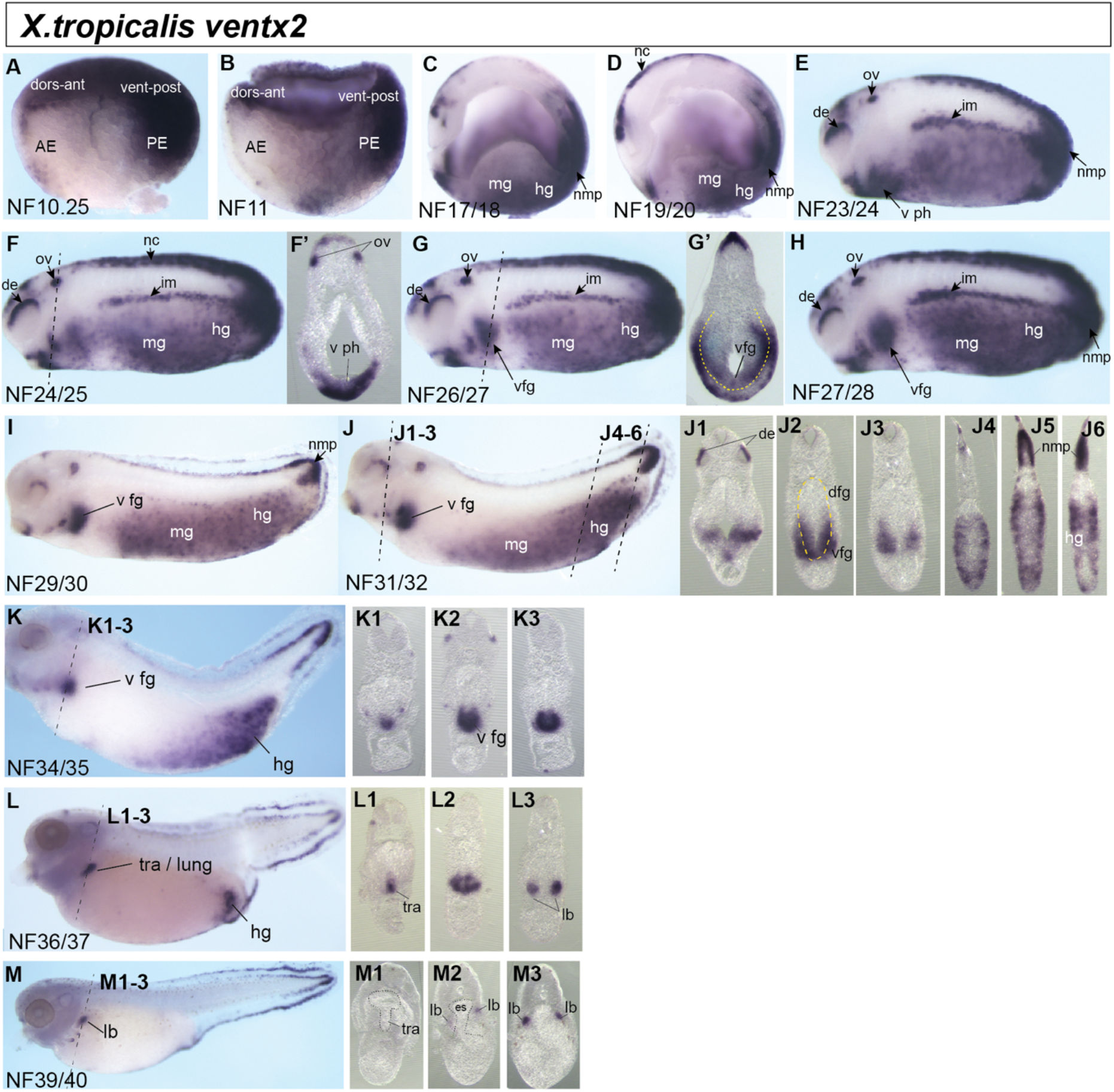
Expression of *ventx2* in developing *Xenopus tropicalis* embryos. (A-M). In-situ hybridization was performed at the indicated stages using a full-length *ventx2*.*1* probe (GenBank sequence CR848273.2), predicted to detect both *ventx2*.*1/ventx2*.*2* due to 92% sequence identity. Abbreviations: AE, anterior endoderm; PE, posterior endoderm; dors-ant, dorsal-anterior; vent-post, ventral-posterior; de, dorsal eye; hg, hindgut; im, intermediate mesoderm; mg, midgut; nmp, neuromesodermal progenitors; nc, neural crest; ov, otic vesicle; dfg, dorsal foregut; v fg, ventral foregut; v ph, ventral pharynx; tra, trachea; lb, lung bud; es, esophagus. Dashed black lines in F,G, and J-M indicate plane of section for those sections shown. Dashed yellow lines in G’ and J2 outline the endoderm layer.

We observed the well-characterized expression of *ventx2* in ventral-posterior endoderm and mesoderm, neural crest (nc), and neuromesodermal progenitor (nmp) domains from gastrula (NF10.5) through neurula stages (NF19/20) (Fig.1A-D). As anterior-posterior axis elongation progressed during NF23-30, *ventx2* was expressed in the dorsal eye (de), otic vesicle (ov), intermediate mesoderm (im), nc, nmps, ventral pharynx (v ph), midgut (mg) and hindgut (hg) regions (Fig.1E-I), with a distinct posterior ventral foregut (v fg) domain of expression becoming evident by NF26-28 (Fig.1G-H, section in Fig1.G’). The ventral foregut expression domain persisted through NF36/37, whereas the midgut and hindgut domains of *ventx2* expression declined during these stages (Fig.1I-L). Sections of the foregut region showed that from NF24-32 *ventx2* was expressed in both the endoderm and mesoderm layers (Fig.1F’G’, J2; yellow dashed lines outline the endoderm), but that after NF34 it was restricted to the ventral foregut endoderm of the trachea/lung forming region (Fig.1K-M).

Importantly the *ventx2* expression in the ventral foregut endoderm at NF31/32 is prior to the onset of *nkx2-1*, currently the earliest known epithelial marker of respiratory progenitor cells, and a temporal comparison of this finding is shown in Fig. 2A. Thus, to our knowledge, *ventx2* is the earliest marker of trachea and lung epithelial progenitors in Xenopus. Interestingly, by NF40 expression was lost from the tracheal domain and only persisted in the distal lung bud tips (Fig1.M) and by NF42 *ventx2* was no longer expressed in any endoderm domains (supplemental Fig.S2).

**Figure 2.**
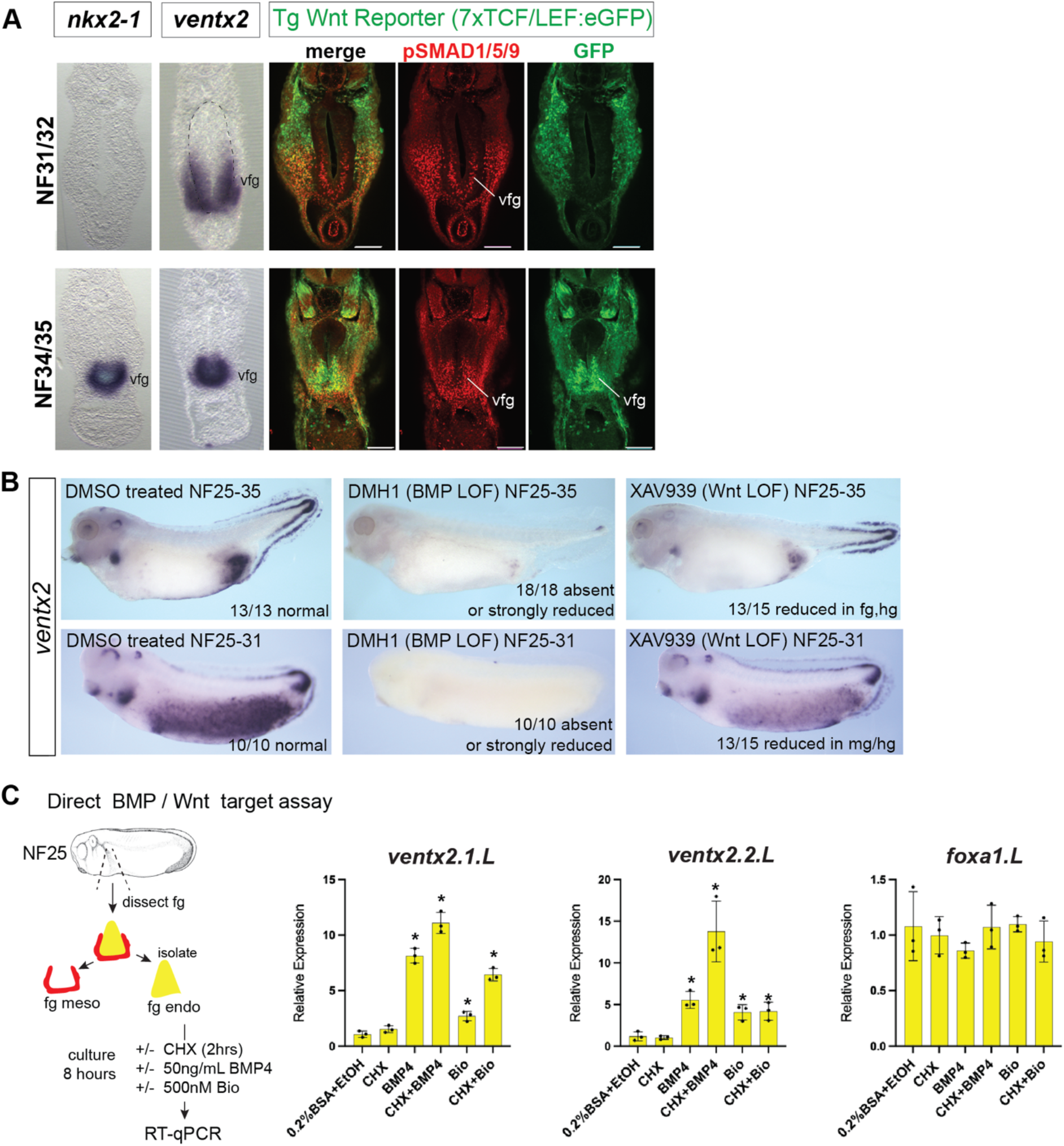
BMP signaling acts prior to Wnt/B-catenin to directly activate *ventx2* expression in foregut endoderm. (A). BMP signaling is active and *ventx2-1* is expressed in respiratory progenitor endoderm prior to *nkx2-1* and Wnt/B-catenin signaling. Comparison of in-situ hybridization for *ventx2* and *nkx2-1* in foregut sections of stage NF31/32 and NF34/35 embryos show expression of *ventx2* in the ventral foregut at NF31/32 prior to onset of *nkx2-1*. Note the *venxt2* sections in this comparison are re-used from figure 1. Immuonstaining of transgenic Wnt/B-catenin reporter *X*.*tropicalis* embryos (transgene carries 7 Tcf/Lef binding sites driving eGFP; Tran et al 2010) with antibodies indicative of active BMP (phosphorylated Smad1/5/9 (pSMAD1/5/9; red) and Wnt/B-catenin (eGFP; green) signaling show that pSMAD1/5/9 is detectable in the ventral foregut endoderm at stage NF31/32 coincident with *ventx2*, whereas active Wnt/B-catenin (eGFP) in the endoderm is detected later at stage NF34/35 coincident with *nkx2-1*. Scale bar=100uM. (B). BMP signaling is required for foregut expression of *ventx2* at NF31/32, whereas both BMP and Wnt/B-catenin are necessary for robust foregut *ventx2* expression at NF35. Embryos were cultured in DMSO (vehicle control) or 20uM DMH-1 (Bmp type I receptor inhibitor) or 25uM XAV939 (Tankyrase inhibitor that triggers B-catenin degradation) from NF25-31or from NF25-35 and assayed by in-situ hybridization for *ventx2*. (C). Direct BMP / Wnt target assay in *X*.*laevis* foregut endoderm explants. Foregut endoderm was isolated at stage NF25 and pre-treated for 2 hours in cycloheximide (CHX) prior to further exposure in CHX with either 50ng/mL BMP4 protein or 500nM Bio and gene expression was assayed by RT-qPCR after 8 hours of total culture. The combination of 0.2% BSA + ethanol (EtOH) is vehicle control treatment. BMP4 robustly induced expression both *ventx2*.*1* and *ventx2*.*2* in the presence of CHX, demonstrating direct activation. *foxa1* is a pan-endoderm gene not significantly affected by BMP or Wnt/B-catenin stimulation. Each black dot in the graphs represents a biological replicate (pool of n=3 explants) and *asterisks=p<0.05, parametric two-tailed T-test relative to vehicle treated explants.

### BMP signaling acts prior to Wnt/B-catenin to directly activate *ventx2* expression in respiratory progenitor cells

As BMP and Wnt/B-catenin are evolutionarily conserved signals that control vertebrate respiratory progenitor induction (Whitsett et al 2019) and *ventx2* is a known BMP and Wnt target in the gastrula (von Bubnoff et al 2005; Hikasa et al 2010), we compared *ventx2* expression in the developing foregut to endogenous BMP signaling, as indicated by immunostaining for phosphorylated Smad1/5/9 (pSmad1), and endogenous Wnt/B-catenin signaling, reflected by eGFP expression, in transgenic Wnt/B-catenin reporter embryos (Fig.2A; Tran et al 2010). At NF31/32, pSmad1 was detectable in the ventral foregut co-incident with *ventx2*, however the respiratory progenitor marker *nkx2-1* and Wnt/B-catenin-dependent eGFP were not yet expressed in the foregut epithelium. This suggests BMP signaling is primarily responsible for the activation of *ventx2* expression in the ventral foregut. By NF34/35, eGFP is detectable in the ventral epithelium, indicating active Wnt/B-catenin signaling, and *nkx2-1* has been induced, reflecting respiratory progenitor induction (Fig.2A).

To test the temporal requirement for BMP and Wnt/B-catenin regulating *ventx2* expression in the ventral foregut, we treated embryos with small molecule inhibitors of each pathway starting at NF25 and assaying embryos at either NF31 or NF35 (Fig.2B). Maximal *ventx2* expression at NF35 was dependent on both pathways, as disrupting BMP signaling with the type I BMP receptor antagonist DMH1 or Wnt/B-catenin with the Tankyrase inhibitor XAV939 each resulted in a loss or severe reduction in foregut expression of *ventx2* (Fig.2B). Importantly, embryos assayed at NF31 revealed that all *ventx2* expression domains, including the ventral foregut, were BMP-dependent whereas XAV939 treatment had little effect on foregut *ventx2* while reducing it in the midgut/hindut (Fig.2B). We also noted the hindgut and tailbud neural crest / neuromesodermal progenitor domains of *ventx2* at NF35 were dependent on BMP but not Wnt/B-catenin signals (Fig.2B). Together these data suggest that BMP signaling in the developing foregut acts prior to Wnt/B-catenin to initiate *ventx2* expression in respiratory progenitor cells and that both pathways ultimately contribute to maximal *ventx2* expression.

To test if BMP and/or Wnt/B-catenin signaling was sufficient to directly activate *ventx2* expression in the foregut epithelium, we performed foregut endoderm explant assays in *X*.*laevis* embryos, which are larger and easier to dissect, stimulating the BMP or Wnt/B-catenin pathways in the presence of cycloheximide (CHX), which blocks protein synthesis and prevents and intermediate factors from being expressed (Fig.2C). We dissected out foreguts from stage NF25 embryos and manually removed the mesoderm/ectoderm layers, which are the endogenous source of Wnt and BMP. Explants were then pre-treated for 2 hours in CHX prior to further exposure in CHX with either 50ng/mL BMP4 protein or 500nM Bio, a small molecule GSK3 inhibitor that stabilizes B-catenin and activates the canonical Wnt pathway, and performed RT-qPCR analysis after 8 hours of total culture. BMP4 treatment robustly activated expression of both *ventx2*.*1* and *ventx2*.*2* in the presence of CHX, demonstrating direct activation (Fig.2C). Bio treatment also upregulated *ventx2* expression in the presence of CHX, albeit to a lesser extent than BMP4 demonstrating that they are also direct Wnt targets. Together these data suggest that both BMP and Wnt/B-catenin can directly contribute to foregut epithelial *ventx2* expression, consistent with the known combinatorial signaling of these pathways driving lung induction.

### Data-mining reveals potential Smad1/B-catenin bound enhancers regulating foregut *ventx2* expression

We next performed data mining of public epigenetic and ChIP-seq data to identify putative BMP and WNT responsive enhancers that might regulate the ventral foregut *ventx2* expression. We searched Xenbase for published *X*.*tropicalis* and *X*.*laevis* ChIP-seq and ATAC-seq data sets at gastrula (NF10-10.5), neurula (NF16, NF20) and tailbud (NF30) stages. Xenbase curates and facilitates the easy downloading of BigWig files that can be viewed with genome browsers such as IgV (Fig.3).

**Figure 3.**
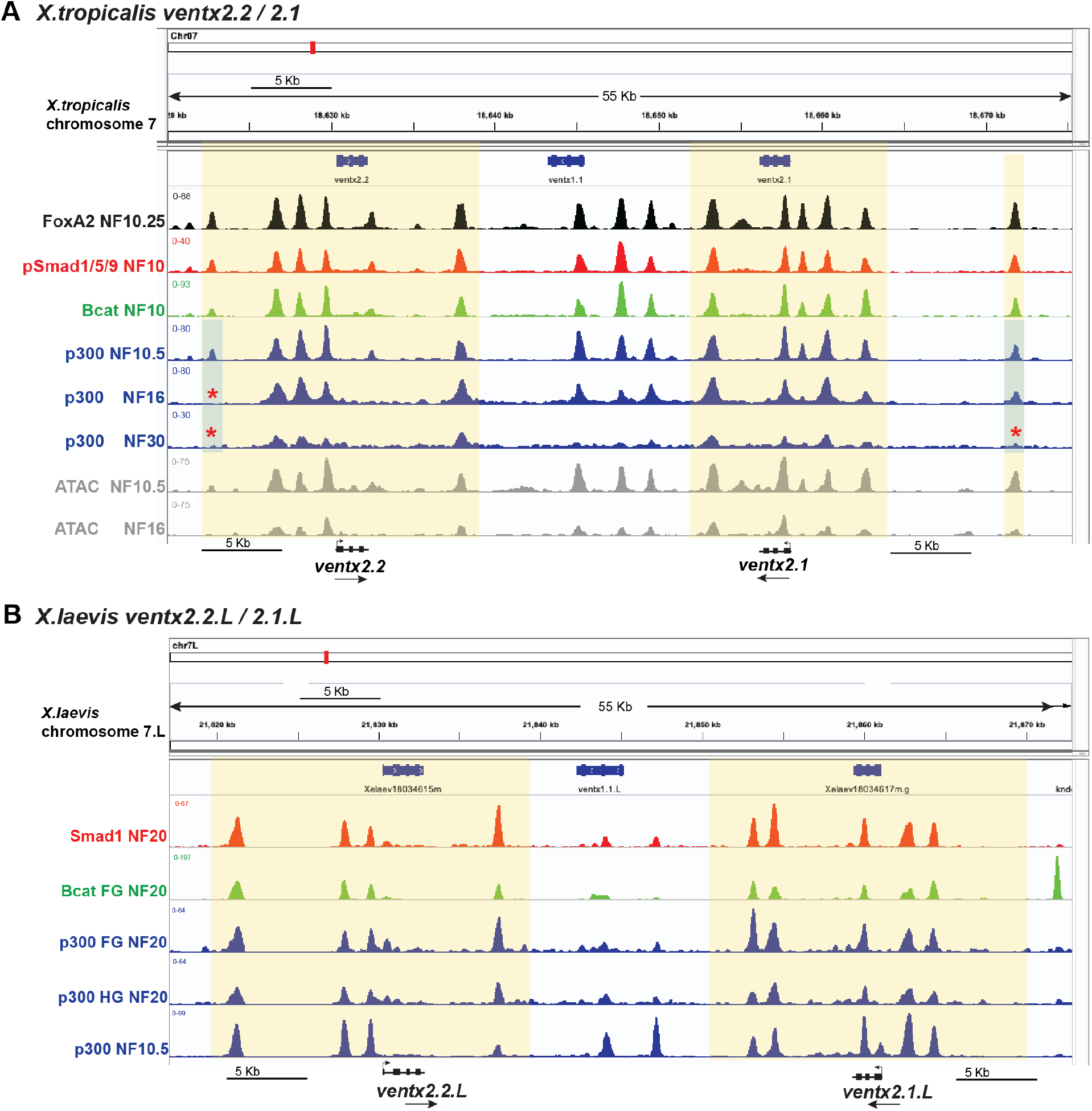
Data mining of published ChIP-seq and ATAC-seq data suggests multiple enhancers around *ventx2*.*1* and *venx2*.*2* are re-utilized during development to regulate *ventx2* expression. (A). IgV browser shot of the genomic locus around *X*.*tropicalis ventx2*.*1/2*.*2* spanning approximately 55 Kilobases (Kb). Tracks display the indicated ChIP-seq or ATAC-seq binding or accessibility data at the indicated NF stage of development. Gene models are drawn in black below and black arrows indicate the 5’-3’ direction of the genes, which lie in opposite orientations. Data were from the following sources: FoxA2 NF10.25: GSE85273 (Charney et al 2017); phosphorylated Smad1/5/9 NF10: GSE113186 (Gentsch et al 2019); B-catenin NF10: GSE113186 (Gentsch et al 2019); p300 NF10.5, NF16, NF30: GSE67974 (Hontelez et al 2015); ATAC-seq NF10.5 and NF16: GSE145619 (Bright et al 2021). Blue shading and red asterisks indicate regions of possible differential p300 binding over developmental time. (B). IgV browser shot of the genomic locus around *X*.*laevis ventx2*.*1/2*.*2* spanning approximately 55 Kb. Tracks display the indicated ChIP-seq data at the indicated NF stage of development. Gene models are drawn in black below and black arrows indicate the 5’-3’ direction of the genes, which lie in opposite orientations. Data were from the following sources: Smad1 NF20: GSE87652 (Stevens et al 2017); B-catenin NF20 foregut explants: GSE87652 (Stevens et al 2017); p300 NF20 foregut and hindgut explants: GSE87652 (Stevens et al 2017); p300 NF10.5: GSE76059 (Session et al 2016).

First we examined the *X*.*tropicalis ventx2*.*1* and *ventx2*.*2* loci during development (Fig.3A). Since gastrula stage *ventx2* expression is known to be directly controlled by BMP/pSmad1 and Wnt/B-catenin signals (von Bubnoff et al 2005; Hikasa et al 2010), we examined gastrula ChIP-seq data for pSmad1 and B-catenin binding (Gentsch et al 2019) along with that for FoxA2, an pioneer endoderm TF useful for identifying endoderm enhancers (Charney et al 2017), p300, a histone acetyltransferase that is typically recruited to and marks enhancers (Hontelez et al 2015), and chromatin accessibility via ATAC-seq (Bright et al 2021). We observed multiple gastrula ChIP-seq peaks upstream and downstream in regions +/-20 Kb from the start of both *ventx2*.*1* and *ventx2*.*2* that were bound by FoxA2, pSMAD1, B-catenin, and p300 which over-lapped with ATAC-seq chromatin accessibility peaks, suggesting a number of enhancers contribute to gastrula *ventx2* expression (Fig.3A). Interestingly, some of the Foxa2/p300/Smad1/B-catenin co-bound peaks from gastrula ChIP-seq were also bound by p300 later in development (NF16-30), suggesting common enhancers regulate both the early and later temporal phases of *ventx2* expression. We do note some minor temporal differences, such as the presence of distal enhancers upstream of both *ventx2*.*1* and *ventx2*.*2* which displayed p300 binding at NF10.5 and/or NF16 but not NF30 (red asterisks/blue shading in Fig.3A; approximately ∼17Kb upstream of *ventx2*.*1* and ∼10Kb upstream of *ventx2*.*2)*, suggesting some degree of putative differential temporal regulation could occur.

We also re-examined our previously published Smad1, B-catenin, and p300 ChIP-seq data on dissected foregut and hindgut tissue from *X*.*laevis* embryos at NF20 (Fig.3B, Stevens et al 2017). Similar to tropicalis, we observed multiple putative enhancers co-bound by Smad1, B-catenin, and p300 in the foregut which are likely to regulate the expression in the presumptive respiratory domain (Fig.3B), While additional ChIP-seq studies are obviously needed in tailbud embryos at NF31 and NF35 during respiratory lineage induction to make definitive conclusions, together these genomic analyses suggest that common enhancers regulating BMP/Smad1 and Wnt/B-catenin dependent gastrula *ventx2* expression are re-utilized to regulate foregut endodermal *ventx2* expression during subsequent development.

### Ventx2 regulates respiratory system development

To investigate the function of Ventx2 in foregut respiratory progenitors, we injected embryos at the 16-cell stage in dorsal-anterior vegetal blastomeres with a previously validated translation blocking morpholino oligo (MO) to knockdown *ventx2*.*1/ventx2*.*2* (Sander et al 2007; Rankin et al 2011; Scerbo et al 2012; Scerbo et al 2017; Maharana and Schlosser 2018; Scerbo et al 2020). Our injection strategy targets the future foregut avoiding the well-characterized gastrula stage axial patterning phenotype that results from depleting Ventx2 function in ventral-posterior mesendoderm (Sander et al 2007; Rankin et al 2011). A representative lineage trace of the targeted foregut cells is shown in Fig.4A. Analysis of Ventx2 foregut morphants at NF42 via in-situ hybridization for the respiratory epithelium differentiation maker *surfactant protein c* (*sftp-c)* revealed a striking increase in lung bud size as well as expanded/ectopic *sftp-c* expression in the anterior pharynx (Fig.4B). Immunostaining and confocal microscopy analysis of the NF42 foregut for Fibronectin, which stains the extracellular matrix (ECM) of the splanchnic mesoderm, and endodermal Foxa2 also revealed a failure of tracheal-esophageal (T-E) separation in Ventx2 foregut morphants (Fig.4C). In addition, in control lung bud sections a lumen is normally present that is lined by Foxa2+ epithelial cells; however a lumen was not evident in Ventx2 foregut morphant lung buds and Foxa2+ cells filled the lung bud space; co-staining with phospho-Histone H3 (pHH3) to identify mitotic nuclei suggested increased numbers of pHH3+ nuclei (Fig.4C). We used Imaris software to generate 3-D renderings of the lung domain from our confocal z-stacks and quantitated the numbers of pHH3+ nuclei within, which revealed a 30.2% average increase in the number of pHH3 positive nuclei in the Ventx2 foregut morphant lung domain compared to controls (n=5 embryos analyzed per condition, Supplemental Fig.S3B; Supplemental Table S1A). Together these data demonstrate Ventx2 regulates lung bud size, is necessary for proper T-E separation, and modulates cell proliferation.

**Figure 4.**
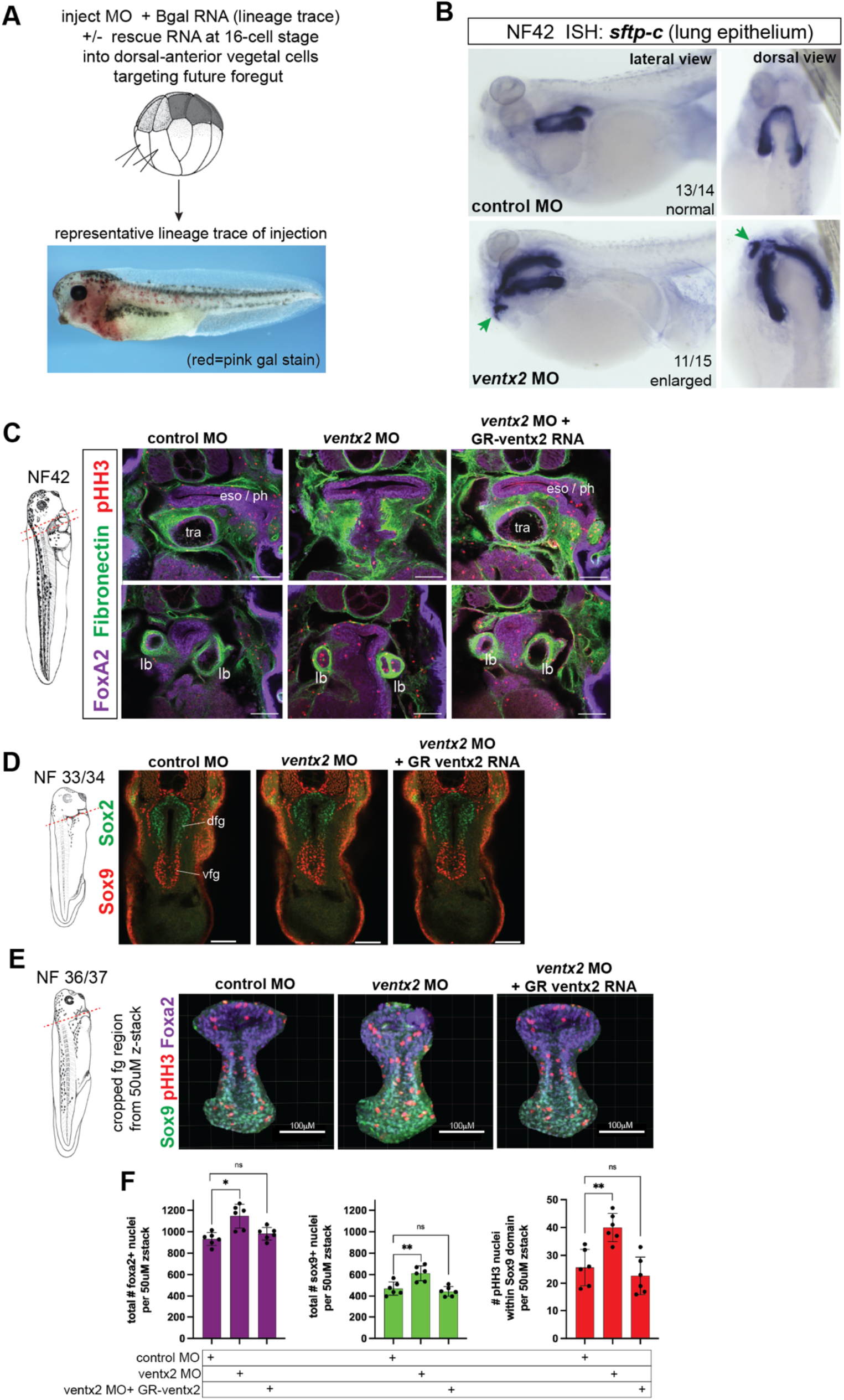
Ventx2 regulates respiratory system development. (A). Microinjection strategy to knockdown Ventx2 in the developing foregut. Embryos at the 16-cell stage were injected in dorsal-anterior vegetal blastomeres to target the future foregut. A representative lineage trace of the injected cells is shown in a NF36/37 embryo via pink-gal staining (red signal) resulting from activity of the injected B-galactosidase mRNA (lineage tracer). (B). In-situ hybridization for the lung epithelium marker *sftp-c* in NF42 embryos (lateral and dorsal views) reveals larger lung buds and expanded *sftp-c* in the pharynx (green arrow) in Ventx2 foregut morphants as compared to control MO injected embryos. The total number of embryos analyzed by *sftp-c* in-situ hybridization that had the observed staining pattern are indicated. (C). Ventx2 is required for tracheal-esophageal separation and lung bud luminal structure. Confocal optical sections of NF42 embryos through the foregut immunostained for the endoderm transcription factor Foxa2 (purple), the splanchnic mesoderm ECM marker Fibronectin (green), and mitotic nuclei marker phospho-Histone H3 (pHH3, red) reveal failed separation of the foregut tube into a distinct dorsal esophagus and ventral trachea in Ventx2 foregut morphants. Abbreviations: eso, esophagus; ph, pharynx; tra, trachea; lb, lung bud. Scale bar=100uM. (D). Dorsal-ventral pattern of the foregut is normal at stage NF33/34 in Ventx2 morphants. 50uM Confocal z-stacks are shown of immunostaining for the ventral respiratory progenitor marker Sox9 (red) and dorsal esophageal progenitor maker Sox2 (green). Abbreviations: dfg, dorsal foregut; vfg, ventral foregut. Scale bar = 100uM. (E-F). Elevated progenitor cell number in the foregut endoderm domain of NF36/37 Ventx2 foregut morphants. Imaris software was used to generate the cropped endoderm foregut region from 50uM confocal z-stacks that were immunostained (E) and quantitated (F) for the numbers of positive nuclei for the pan-foregut endoderm transcription factor Foxa2 (purple), ventral respiratory progenitor marker Sox9 (green) and mitotic marker phospho-Histone H3 (pHH3, red); scale bar = 100uM. Each black dot in the graphs in (F) is a separate embryo 50uM cropped endoderm confocal z-stack (n=6 quantitated per condition). Asterisks *=p<0.05,**=p<0.01, parametric two-tailed T-test; ns=not significant.

**Figure 5.**
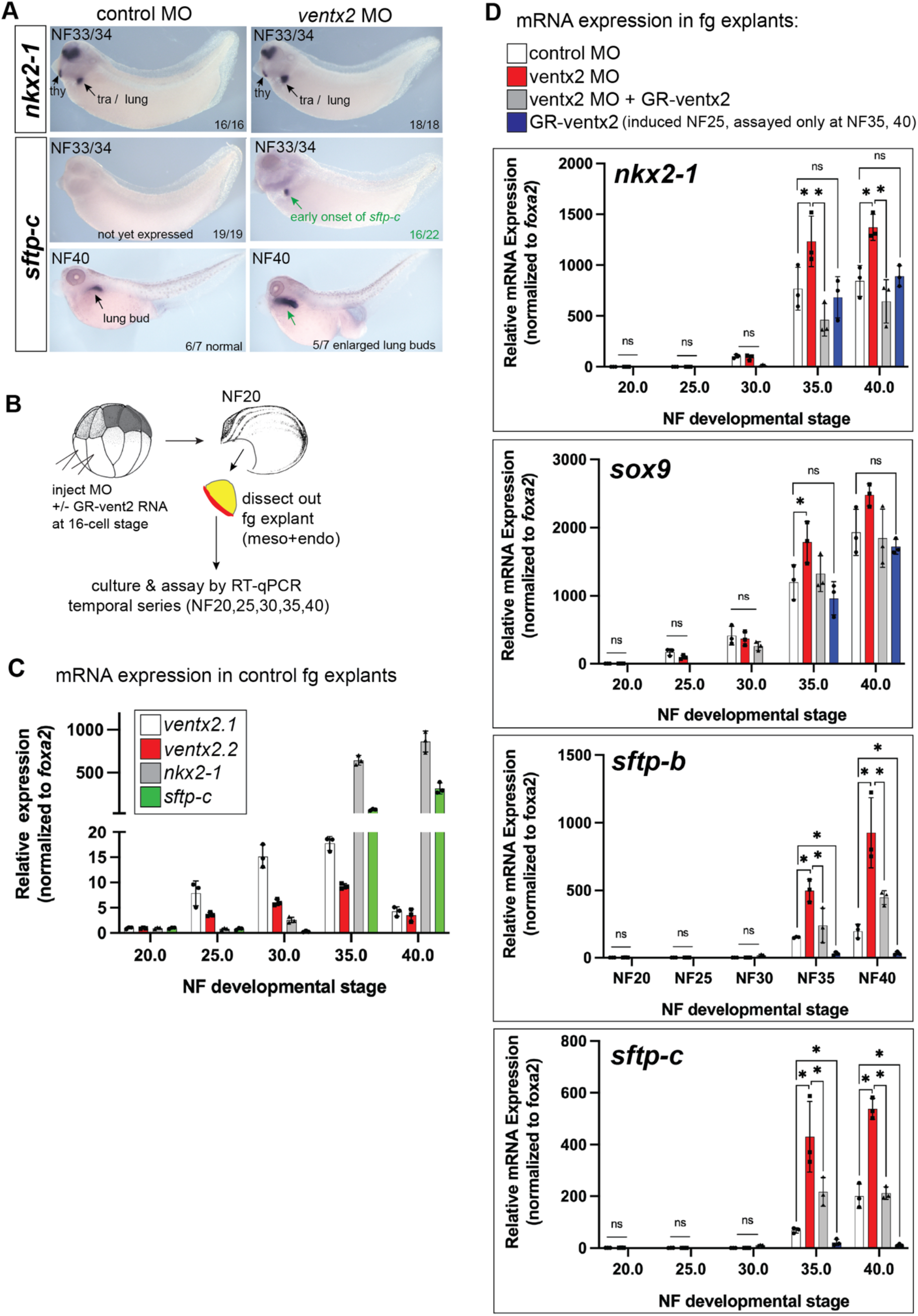
Ventx2 regulates the timing and level of surfactant protein gene expression. (A). Expression of the lung epithelial marker *sftp-c* was precociously activated at NF33/34 in Ventx2 foregut morphants, which have larger lung buds at NF40. In situ hybridization of *nkx2-1* and *sftp-c* at NF33/34 and NF40. (B). Schematic of foregut explant and culture assay to assess temporal kinetics of respiratory marker gene expression by RT-qPCR. (C). RT-qPCR analysis of control MO injected foregut explants demonstrates that gene expression in the fg explant culture system mirrors normal embryonic development, with ventx2.1 and ventx2.2 expression (white and red bars) increasing from NF25-35, but is lower at NF40. *sftp-c* expression (green bars) is not detectable prior to NF35 and follows *nkx2-1* (gray bars). (D). RT-qPCR analysis of control, Ventx2 loss of function, rescued, or gain of function fg explants. Progenitor markers *nkx2-1* and sox9 were not prematurely activated from NF20-30 in response to Ventx2 depletion, however differentiation markers *sftp-b* and *sftp-c* were prematurely elevated. Ventx2 gain of function (blue bars; GR-venxt2 construct was Dex-activated at NF25 and gene expression was assayed at NF35 and NF40) resulted in suppression of *sftp-b* and *sftp-c* respiratory differentiation markers but had no effect on the respiratory progenitor markers *nkx2-1* or *sox9*. Each black dot in the graphs in C,D represents a biological replicate (pool of n=3 explants) and *asterisks=p<0.05, parametric two-tailed T-test relative to uninjected stage NF20 fg explants.

To determine if the larger lung buds in Ventx2 morphants were associated with increased numbers of respiratory progenitor cells earlier in development, and to determine if the disrupted T-E separation was associated with earlier foregut patterning defects, we analyzed embryos and earlier timepoints, at NF33/34 and NF36/37 via immunostaining and confocal microscopy (Fig.4D-F). At NF33/34, no obvious dorsal-ventral pattern defect was obvious in Ventx2 foregut morphants, as assessed by immunostaining for Sox2, which marks dorsal esophageal, and Sox9 which marks ventral respiratory progenitors, as their normal dorsal-ventral pattern of localization was normal (Fig.4D). However quantitation of the numbers of Sox2+ and Sox9+ nuclei in 50uM foregut confocal z-stacks did reveal an 26% average increase in in Sox9+ nuclei in the Ventx2-depleted foregut at NF33/34 but no difference in Sox2+ nuclei number (n=5 embryos analyzed per condition, Supplemental Fig.S3A; Supplemental Table S1B).

Additional immunostaining, confocal analysis, and quantitation of 50uM foregut z-stacks at NF36/37 for Foxa2 / pHH3 / and Sox9 (Fig4.E,F) again showed normal dorsal-ventral pattern at this stage but a 30% average increase in the number of Sox9+ nuclei in the ventral foregut of Ventx2 morphants compared to controls (n=5 embryo foregut domains analyzed per condition; Fig.4F, Supplemental Table S1C). We further quantified the number of pHH3+ mitotic nuclei within the respiratory Sox9 domain and observed a 56% average increase in pHH3+ nuclei just within the Sox9 domain of Ventx2 morphants (Fig.4F; Supplemental Table S1D). Together these data suggest Ventx2 restrains proliferation of foregut endoderm progenitors during the earliest stages of lung development in Xenopus.

### Ventx2 regulates the timing and level of surfactant protein gene expression

Since Ventx2 has previously been shown to suppress premature differentiation in the blastula stage ventral mesendoderm and in the developing neural crest (Scerbo et al 2012; Scerbo et al 2020), we next investigated the timing and expression level of surfactant protein encoding genes *sftp-b* and *sftp-c*, which mark respiratory epithelial differentiation (Hyatt et al 2007) and are activated by the master pulmonary TF Nkx2-1 (Tagne et al 2012; Kuwahara et al 2020). In Xenopus *nkx2-1* is first detectable by in-situ hybridization at NF33/34 whereas *sftp-c* onset occurs approximately 7-8 hours later at NF35/36. In situ hybridization analysis of Ventx2 fg morphants compared to controls revealed an earlier onset of *sftp-c* expression at NF33/34 (Fig.4A), co-incident with the timing of *nkx2-1* expression. To more quantitatively investigate the temporal kinetics of gene expression when Ventx2 is depleted, we performed a RT-qPCR time course analysis of control and ventx2-MO injected *X*.*laevis* foregut (fg) explants. Fg explants containing both the BMP-producing foregut mesoderm and recipient endoderm were dissected at NF20 and cultured for 48 hours to a developmental time when control sibling embryos were at stage NF40; explants were harvested during the culture period at NF25, 30, 35 and 40 (Fig.4B). Expression of respiratory markers *nkx2-1, sox9, sftp-b* and *sftp-c* was determined by RT-qPCR and expression was normalized to the pan-foregut TF *foxa2* to account for changing endoderm cell numbers over time. We first verified that gene expression in these fg explants mirrored normal embryonic development: expression of *ventx2*.*1/2*.*2* increased at NF30 and NF35 but had lower levels at NF40; as expected expression of the respiratory epithelial progenitor markers *nkx2-1* and *sox9* was robust at NF35, and *sftp-c* expression was first detectable at NF35 and increased at NF40 (Fig.4C,D). Thus, the kinetics of respiratory lineage development in these fg explants mirrored normal development.

Interestingly, the temporal onset of *nkx2-1* and *sox9* expression (progenitor markers) was not affected by Ventx2-depletion, as they were not elevated at NF20, NF25, nor NF30; however ventx2-MO embryos did have higher *nkx2-1* and *sox9* expression at NF35 and NF40 (Fig.4D). Consistent with the in situ hybridization data for *sftp-c*, ventx2-MO explants exhibited elevated expression of *sftp-b* and *sftp-c* at NF35 but interestingly not at NF20, NF25, or NF30 (Fig4D). The fact that we did not detect ectopic elevation of these respiratory differentiation genes prior to the time when *nkx2-1* is normally expressed is consistent with previous studies demonstrating that activation of *sftp* transcription is directly dependent on the master pulmonary TF Nkx2-1 binding their promoter/enhancer elements (Kelly et al 1996; Glasser et al 2000; Tagne et al 2012; Little et al 2019; Kuwahara et al 2020). Together these data suggest Ventx2 regulates differentiation timing and *sftp* levels in the respiratory lineage, likely in an Nkx2-1 dependent manner.

We further tested if overexpression of Ventx2 could rescue the MO phenotype or repress *sftp* expression, injecting an dexamethasone (DEX) inducible GR-ventx2 construct (McLin et al 2007). Indeed when GR-ventx2 was co-injected and DEX activated at NF25, this reduced the elevated *nkx2-1, sox9*, and *sftp* expression levels observed in Ventx2 fg morphant explants back to near control levels (Fig4.D; gray bars). Intriguingly, in a gain of function setting, where we induced GR-ventx2 in control explants at NF25, Ventx2 was sufficient to suppress expression of the differentiation indicators *sftp-b* and *sftp-c* but had no effect on the progenitor markers *nkx2-1* or *sox9* (Fig.4D; blue bars). These data further suggest Ventx2 can regulate differentiation timing and levels of *sftp* expression in the developing respiratory lineage, perhaps acting in parallel or downstream of Nkx2-1.

## Discussion

Our data show that *ventx2* is a useful spatial-temporal marker of Xenopus ventral foregut respiratory epithelial progenitors prior to *nkx2-1*, that its early expression is directly induced by BMP signaling, and temporally *ventx2* expression declines as respiratory epithelium differentiates (summarized in Fig.6A,B). Our data mining and analyses of published genomic studies in Xenopus indicate the possible recurrent use of proximal enhancers around *ventx2*.*1/2*.*2* by BMP/pSmad1 and Wnt/B-catenin during multiple stages of development. BMP signaling can activate expression of human *VENTX* during the directed differentiation of human embryonic stem cells into mesoderm and Smad1 binding motifs have been identified in the human VENTX promoter region (Sumi et al 2008; von Bubnoff et al 2005). In the future it will be of interest to perform pSmad1 and B-catenin ChIP-seq during respiratory lineage induction from NF31-35 on isolated Xenopus foregut endoderm tissue to unambiguously define the genomic landscape of BMP and Wnt regulated enhancers at these key developmental stages, and couple this with RNA-seq and functional enhancer studies that are readily performed in Xenopus (Blitz and Cho 2021; Rankin et al 2021).

**Figure 6.**
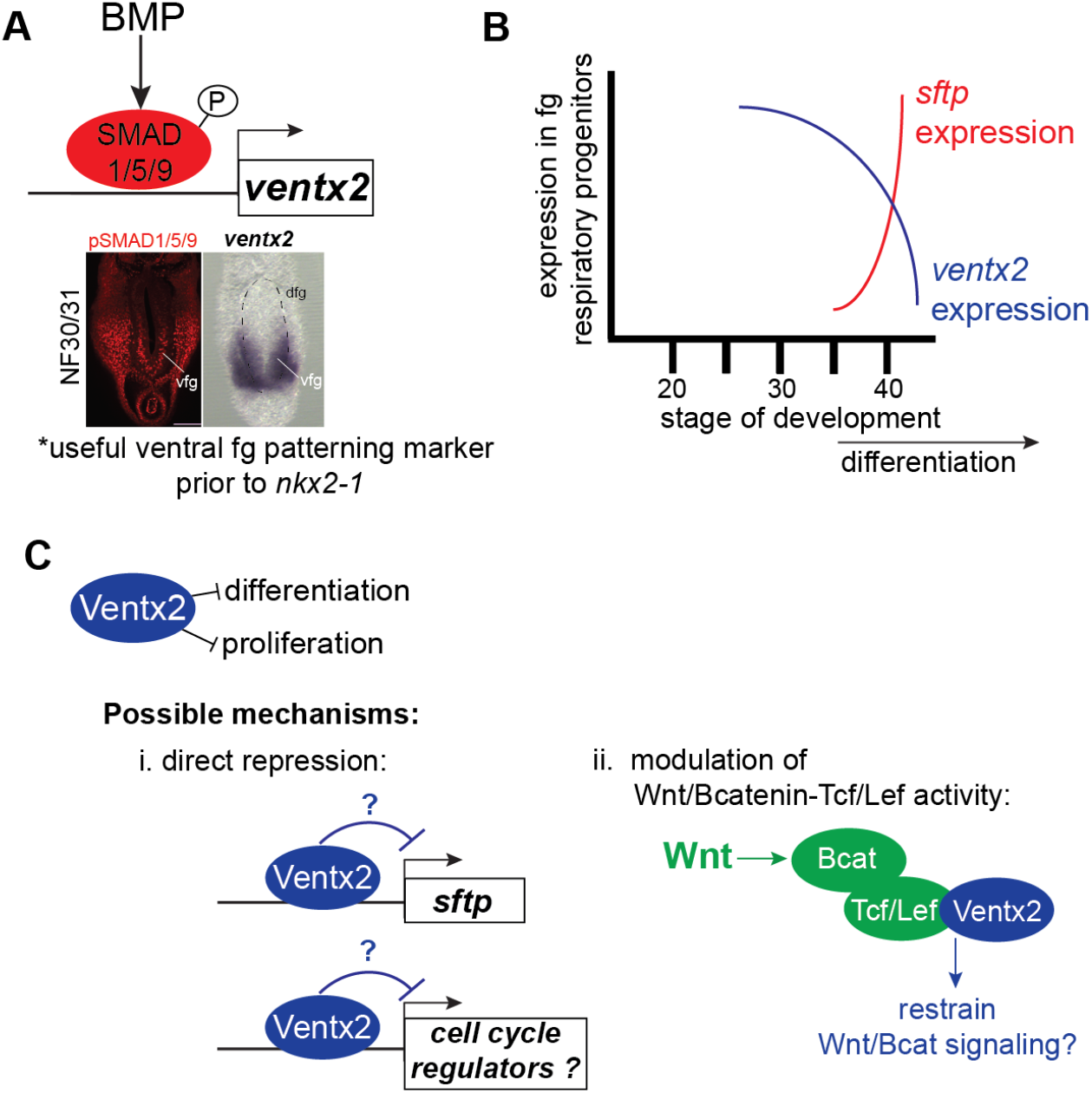
Summary of *ventx2* expression and possible mechanistic functions in developing Xenopus respiratory progenitors.

The phenotypes we observe when depleting Ventx2 in the developing foregut include elevated proliferation, elevated numbers of respiratory progenitors, premature onset and elevated levels of surfactant protein gene expression, and a failure of proper T-E separation. It is important to note that in many contexts premature differentiation of progenitor cells is either associated with or has been shown to be a causative driving force of a myriad of developmental defects or disease states in numerous organ systems including the heart, lung, kidney, and brain (Rowton et al 2021; Morrisey 2013; Chung et al 2018; Kopan et al 2014; Nakafuku and Del Águila 2020; Caracci et al 2021; Khanbabaei et al 2019). In both Xenopus and mice, there is a brief temporal delay between respiratory progenitor induction and the onset of respiratory epithelial differentiation. In Xenopus, *nkx2-1* is first detectable by in-situ hybridization at NF33/34 whereas *sftp-c* onset approximately 7-8 hours later at NF35/36. In mice, *Nkx2-1* is first detectable in the ventral foregut around the 17 somite stage at embryonic day E9.25 (Havrilak & Shannon 2017) while *Sftp-c* onset occurs approximately a day later at E10-10.5 (Wert et al 1993; Ikonomou et al 2017). While rodents have lost the *ventx* gene family during evolution (Kumar et al 2022), it is an intriguing possibility that a conserved BMP-dependent mechanism still controls timing of respiratory epithelial differentiation in vertebrates, perhaps ensuring progenitor number and gene expression programs of foregut endoderm cells are compatible with subsequent T-E morphogenesis. Indeed, in mice BMP is known to act prior to Wnt/B-catenin to suppress Sox2 expression in the ventral foregut to promote respiratory fate induction and proper levels of both BMP and Wnt signals are necessary for T-E separation (Domyan et al 2011; Woo et al 2011; Billmyre et al 2015). It is an open question if the premature differentiation (precocious and elevated expression of *sftp-b/c*) is related to or drives the defective T-E separation we observe in Xenopus Ventx2 foregut morphants. We have recently characterized evolutionarily conserved cell biological and morphogenetic hallmarks that occur during T-E separation (Nasr et al 2019; Edwards et al 2021), and in the future it will be of interest to investigate if these events are disrupted in response to Ventx2 depletion as well as in response to temporal BMP and Wnt pathway loss or gain of function.

Previous studies have identified settings in which Vent factors modulate Wnt/B-catenin signaling, having either potentiating or dampening effects depending on the context (Gao et al 2007; Wu et al 2018). The Ventx2 foregut loss-of-function phenotype we observed of ectopic *sftp-c* is similar to temporal Wnt/B-catenin gain-of-function in the developing foregut (Rankin et al 2012; Rankin et al 2015) and is consistent with the possibility that Ventx2 physically interacts with Tcf/Lef factors to restrain Wnt/B-catenin signaling during early respiratory system development and T-E separation (Fig.6C). Physical interaction between Xenopus Ventx2.2 and Tcf/Lef in the gastrula embryo potentiated Wnt/B-catenin signaling (Gao et al 2007); however in a different context Ventx2.2 regulated mRNA expression levels of *gsk3* and consequently triggered the Gsk3-dependent proteolysis of B-catenin (Wu et al 2018). Physical interaction between human VENTX and human TCF/LEF has also been demonstrated and was associated with suppression of Wnt/B-catenin signaling and suppression of cell proliferation (Gao et al 2010), consistent with our observations on pHH3+ mitotic cell numbers in foregut Ventx2-depleted Xenopus embryos. Together these previously published observations suggest that elevated Wnt/B-catenin activity in the foregut of Ventx2 morphants could partially explain the phenotypes we observe regarding the numbers of respiratory progenitors, cell proliferation, and T-E separation; this possibility warrants further investigation in the future.

In the early blastula Xenopus embryo, Ventx2 has been shown to have a similar function as mouse Nanog and restrains differentiation (Scerbo et al 2012, Scerbo et al 2017). Nanog is known to regulate the pluripotency status of both mouse and human embryonic stem cells and the expression of a number of cell cycle regulators are known to be targets of Nanog (Zhang et al 2009; Liang et al 2008). In general, Ventx2 is a transcriptional repressor (reviewed in Kumar et al 2022), and it is possible Ventx2 could directly repress either *sftp* or cell cycle genes in developing Xenopus respiratory progenitors (Fig.6C).

Interestingly, during mouse lung development BMP signaling is required to maintain the undifferentiated state of airway and alveolar epithelial progenitors, preventing premature exhaustion of their pools and securing the continuous supply of the differentiated epithelial cells necessary for a fully developed lung (Sountoulidis et al 2012). During mouse lung injury studies, reactivation of BMP signaling is observed in injured epithelium with similar requirements regarding proper management of the undifferentiated epithelial progenitor pools are suggested to be necessary (Sountoulidis et al 2012). Examination of the Human Protein Atlas for single-cell RNA-seq *VENTX* expression in human organs (https://www.proteinatlas.org/ENSG00000151650-VENTX/single+cell+type) intriguingly indicates *VENTX is* expressed at a low level in respiratory epithelial alveolar type II (AT2) cells. BMP signaling is active AT2 cells and Niche-mediated BMP/pSmad1 signaling regulates lung alveolar stem cell proliferation and differentiation (Chung et al 2018). Wnt/B-catenin signaling also drives AT2 cell identity and promotes AT2 cell proliferation (Frank et al 2016; Zacharias et al 2018; Nabhan et al 2018; Xi et al 2019). It is tempting to speculate that some of the functions Ventx2 performs in Xenopus embryonic respiratory progenitors modulating proliferation and differentiation, downstream of BMP and Wnt signaling, are evolutionarily conserved and also performed by human VENTX, downstream of BMP and Wnt, in human AT2 respiratory progenitor cells during either normal homeostasis or in response to injury.

## Materials and Methods

### Ethics Statement

Xenopus experiments were performed according to Cincinnati Children’ Hospital Instititutional Animal Care and Use Committee (IACUC) protocol CCHMC 2019-0053.

### Xenopus embryo methods

Ovulation, in-vitro fertilization and natural mating, embryo de-jellying, and microinjection were performed as described (Sive et al., 2000; Lane and Khokha 2022). Embryos were staged by the Nieuwkoop and Faber (NF) normal table of *Xenopus laevis* development (Nieuwkoop and Faber 1994). Wild-type adult *X. laevis* and *X. tropicalis* frogs were purchased from Nasco (Fort Atkinson, WI) or Xenopus1 Corp. (Dexter, MI). Adult transgenic *X. tropicalis* Wnt/B-catenin reporter frogs (*Xtr*.*Tg(WntREs:dEGFP)*^*Vlemx*^, NXR_1094) were purchased from the National Xenopus Resource (Woods Hole, MA; RRID:SCR_013713). The previously validated translation-blocking morpholino oligo (MO) against *ventx2*.*1/2*.*2* (Sander et al 2007; Rankin et al 2011; Scerbo et al 2012; Scerbo et al 2017; Maharana and Schlosser 2018; Scerbo et al 2020) was purchased from GeneTools (Philmath, OR) and injected at the 16-cell stage into embryos with clear dorsal-ventral pigment differences into dorsal-vegetal blastomeres to target the foregut. The ventx2 MO 5′-GTCATCTTGTCTGTATTAGTCCT-3′ or control MO (mismatch MO that differed by 5 base pairs indicated in <u>bold</u>: 5’-<u>A</u>TCA<u>G</u>CTT<u>A</u>TCT<u>A</u>TATTAGT<u>A</u>CT-3’) were injected at 6.75ng/blastomere (13.5ng total). The inducible pT7Ts-HA-GR-ventx2 plasmid construct was previously generated (McLin et al 2007) and the pCS2+nucB-galactosidase plasmid was a gift from Olga Ossipova and Sergei Sokol. Linearized plasmid templates were used to make mRNA for injection using the Ambion mMessage mMachine SP6 or T7 RNA Synthesis kits (ThermoFisher AM1340, AM1344). Total amounts of injected mRNA were as follows: GR-ventx2 RNA, 400pg (200pg per blastomere); B-gal 125pg (62.5pg per blastomere). 1 μM dexamethasone (DEX; Sigma D4902) was used to induce the GR-ventx2 construct. Pink/Red-Gal substrate for B-gal activity detection (lineage tracing) was purchased from Biotium (10012).

### In-situ hybridization

In-situ hybridization of *Xenopus* embryos was performed as described (Sive et al., 2000) with minor modifications. Briefly, embryos were fixed overnight at 4°C in MEMFA (0.1 M MOPS, 2 mM EGTA, 1 mM MgSO4, and 3.7% formaldehyde), washed 3× 5 min in MEMFA without formaldehyde, bisected in this solution (gastrula-NF20), then dehydrated directly into 100% ethanol, washed 5–6 times in 100% ethanol, and stored at −20°C for at least 24 hr. Proteinase K (PK) (ThermoFisher AM2548) on day 1 was used at 2 µg/ml for 10 min on embryos up to stage NF20; on older embryos 7.5 µg/ml PK was used for 15 minutes; hybridization buffer included 0.1% SDS; RNAse A (ThermoFisher 12091021) used at 0.5 µg/ml; and anti-DIG-alkaline phosphatase antibody (Sigma 11093274910) used at 1:5,000 in MAB buffer (100 mM Maleic acid, 150 mM NaCl, and pH 7.5) + 20% heat-inactivated lamb serum (Gibco 16070096) + 2% blocking reagent (Sigma 11096176001). Anti-sense DIG-labeled in-situ probes were generated using linearized plasmid cDNA templates with 10X DIG RNA labeling mix (Sigma 11277073910) according to the manufacturer’s instructions. The tropicalis ventx2.1 cDNA clone used to prepare the ventx2 in-situ probe was picked from our in-house *X*.*tropicalis* neurula cDNA library (clone TNeu125n01); we sequenced the clone from both ends and verified it is identical to GenBank sequence CR848273.2.

### Immunofluorescence

Embryos were fixed in 100 mM HEPES (pH 7.5), 100 mM NaCl, 2.7% methanol-free formaldehyde for 2 hours at room temperature, dehydrated directly into ice-cold Dent’s post-fixative (80% Methanol / 20% DMSO), washed five times in Dent’s, and stored in 100% methanol at −20°C for at least 48 hours. Embryos were serially rehydrated as follows: first into Dent’s (10 minutes at room temperature), then into Dent’s Bleach (80% methanol, 10% DMSO, 10% hydrogen peroxide; incubated in Dent’s bleach for 1 hour at room temperature), then for 10 minutes per wash in a graded methanol series (75%, 40%, 25%), and finally into PBS +0.1% TritonX-100 (PBSTr). Embryos were then cut in a transverse plane through the pharynx and posterior to the liver to create a foregut sample using a fine razor blade on a 2% agarose-coated dish in PBSTr. Foreguts were subjected to antigen retrieval in 1× R-universal epitope recovery buffer (Electron Microscopy Sciences 62719-10) for 1 hour at 60–65°C, washed 2×10 min in PBSTr, blocked for 1–2 hours in PBSTr +10% normal donkey serum (Jackson ImmunoResearch 017-000-001) + 1% DMSO at room temperature, and incubated overnight at 4°C in this blocking solution + primary antibodies. The following primary antibodies were used:

goat anti-Foxa2 (Santa Cruz Biotechnology sc-6554x; diluted 1:1,000); mouse anti-Sox2 (Abcam ab79351; 1:1,000); rabbit anti-Sox9 (EMD Millipore AB5535; 1:1,000); mouse anti-Sox9 (Abcam ab76997, 1:1,000); mouse anti-Fibronectin 4H2 (Developmental Studies Hybridoma Bank 4H2-c, 1:1,000); rabbit anti phospho-Histone H3 (Cell Signaling Technology #53348, 1:1,000); rabbit anti-phosopho-Smad1/5/9 (Cell Signaling Technology #13820, 1:500); chicken anti-GFP (Aves Labs GFP-1020, 1:1,000).

Secondary antibodies included donkey anti-mouse 488, donkey anti-chicken 488, donkey anti-rabbit Cy3, and donkey anti-mouse Cy5, donkey anti-goat 405 (Jackson ImmunoResearch 715-546-151, 703-546-155, 711-166-152, 715-175-151, and 705-476-147, respectively; all used at 1:1,000 dilution). After extensive washing in PBSTr, samples were incubated overnight at 4°C in PBSTr +1% DMSO+secondary antibodies.

Samples were again extensively washed in PBSTr, dehydrated into 100% methanol, washed five times in 100% methanol, cleared, and imaged in Murray’s Clear (two parts benzyl benzoate, one part benzyl alcohol) on a metal slide with glass coverslip bottom using a Nikon A1R confocal microscope to obtain optical sections.

### Foregut explant dissections and RT-qPCR

Foregut explants were micro-dissected in 1×MBS (Sive et al 2000) + 50 μg/mL gentamycin sulfate (gent; MP Biochemicals 1676045) ± 10 U/mL dispase (Corning Life Sciences 354235; to help remove the mesoderm/ectoderm layers where applicable) and were cultured in 0.5× MBS +0.2% fatty acid free BSA (Fisher BP9704) + 50 μg/ml gent with the following concentrations of factors: 1 μM cycloheximide (CHX; Sigma C1988); 50 ng/ml BMP4 (R&D Systems 314-BP-010); 500nM Bio (Cayman Chemical 13123). In CHX experiments, explants were treated for 2 hours in CHX prior to treatment in CHX+BMP4 or Bio for 6 hours. In the time course analysis in Fig.4C-D, sibling embryos were cultured along in parallel, and explants were frozen at the corresponding stage of the sibling embryo’s developmental age as judged by the Nieuwkoop and Faber (NF) normal table of *Xenopus laevis* development (Nieuwkoop and Faber 1994).

Explants were frozen on dry ice in 200 μl of TRIzol (ThermoFisher 15596018) and stored at –80°C. RNA was extracted using TRIzol and purified using the Direct-zol RNA miniprep plus kit (ZymoResearch R2070); 500 ng RNA was used in cDNA synthesis reactions using Superscript Vilo Mastermix (ThermoFisher 11755050), all according to the manufacturer’s instructions. qPCR reactions were carried out using PowerUp Mastermix (ThermoFisher A25742) on an ABI QuantStudio3 qPCR machine. *Xenopus* RT-qPCR primer sequences are listed in supplementary table S2. Relative expression, normalized to ubiquitously expressed gene *odc* and/or to the foregut endoderm TF *foxa2*, was determined using the 2^−ΔΔCt^ method. Graphs display the average 2^−ΔΔCt^value ± standard deviation. Statistical significance (p<0.05) was determined using parametric two-tailed paired t-test, relative to uninjected, untreated explants. Each black dot in the RT-qPCR graphs represents an independent biological replicate containing n=3 explants.

### Data mining of published ChIP-seq and ATAC-seq data

Bigwig files for data sets listed below were downloaded from Xenbase (http://www.xenbase.org/, RRID:SCR_003280) and viewed using the Integrated Genomics Viewer version IgV_2.5.2 browser, freely downloaded from the Broad Institute (https://software.broadinstitute.org/software/igv/).

Published *X*.*tropicalis* ChIP-seq and ATAC-seq data sets utilized include: FoxA2 NF10.25: GSE85273 (Charney et al 2017); phosphoSmad1/5/9 NF10: GSE113186 (Gentsch et al 2019); B-catenin NF10: GSE113186 (Gentsch et al 2019); p300 NF10.5, NF16, NF30: GSE67974 (Hontelez et al 2015); ATAC-seq NF10.5 and NF16: GSE145619 (Bright et al 2021).

Published *X*.*laevis* ChIP-seq data sets utilized include: Smad1 NF20: GSE87652 (Stevens et al 2017); B-catenin NF20 foregut explants: GSE87652 (Stevens et al 2017); p300 NF20 foregut and hindgut explants: GSE87652 (Stevens et al 2017); p300 NF10.5: GSE76059 (Session et al 2016).

## COMPETING INTERESTS

No competing interests declared.

## FUNDING

This work was funded by NIH Grants P01 HD093363 and 1R01DK123092 to A.M.Z.

## Figures and Figure Legends

**Supplemental Figure S1.**
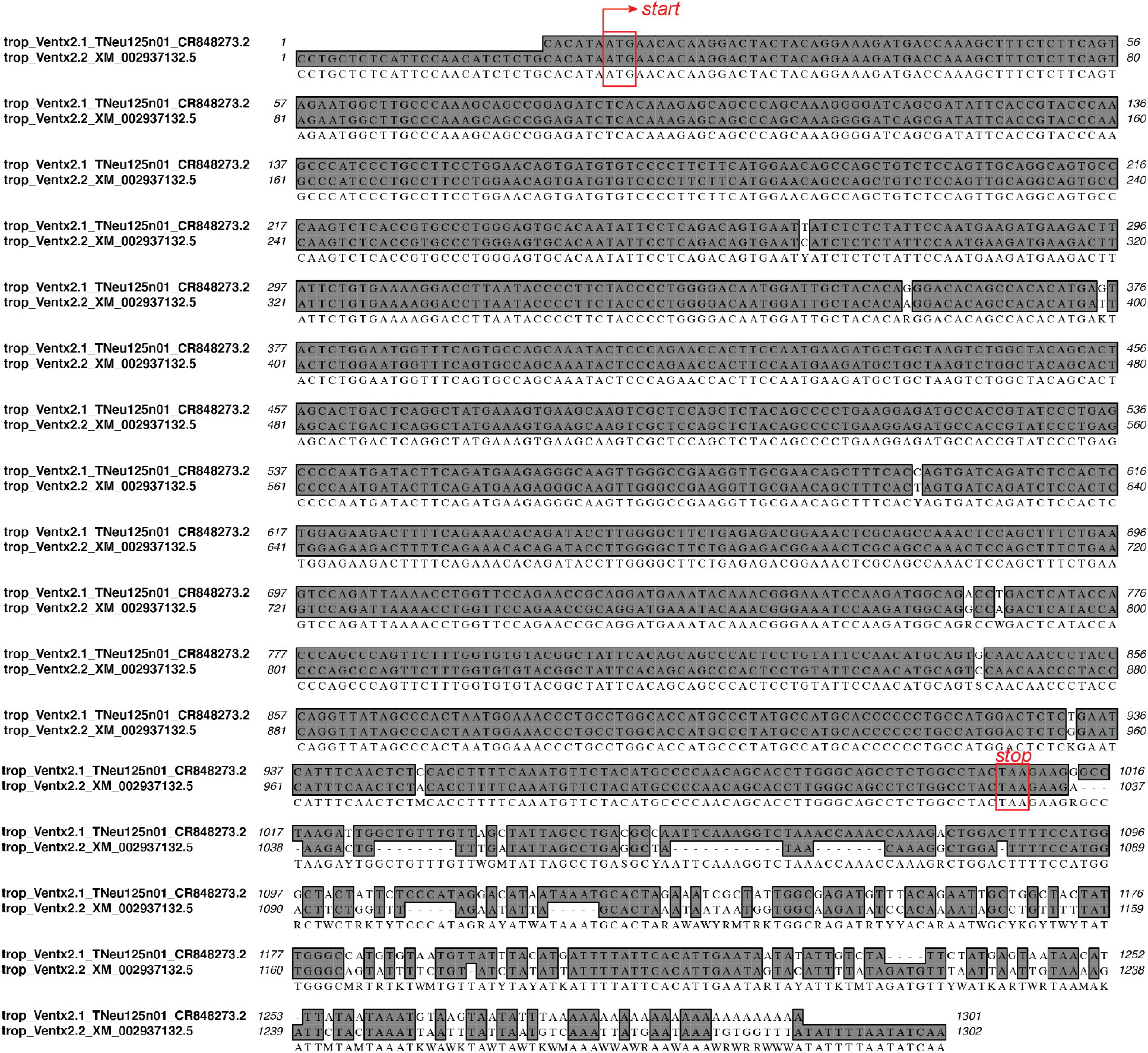
Sequence alignment of *X*.*tropicalis ventx2*.*1*/*ventx2*.*2*. Clustal alignment comparing *X*.*tropicalis* 1301 base pair (bp) *ventx2*.*1* GenBank sequence CR848273.2 to 1302 bp *ventx2*.*2* NCBI reference sequence XM_002937132.5. Nucleotides encoding the start and stop codons are boxed in red. Gray shading indicates identical nucleotides.

**Supplemental Figure S2.**
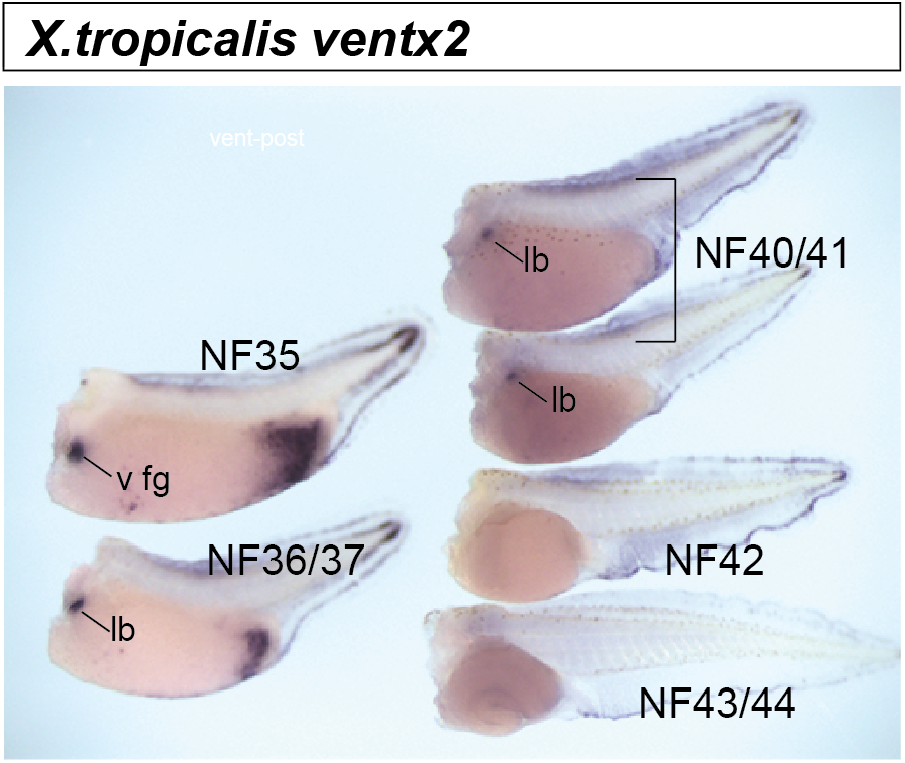
*ventx2* expression is not maintained as development progresses. In-situ hybridization for *ventx2* at the indicated stages. To ensure probe penetration, the heads of the embryos were cut off after fixation and embryos were processed simultaneously in a single vial during the complete in-situ hybridization procedure. *ventx2* is detected in the ventral foregut (v fg) and lung bud (lb) and NF35 and NF36/37, weakly detected at NF40/41, and not detected in NF42 or NF43/44 embryos.

**Supplemental Figure S3.**
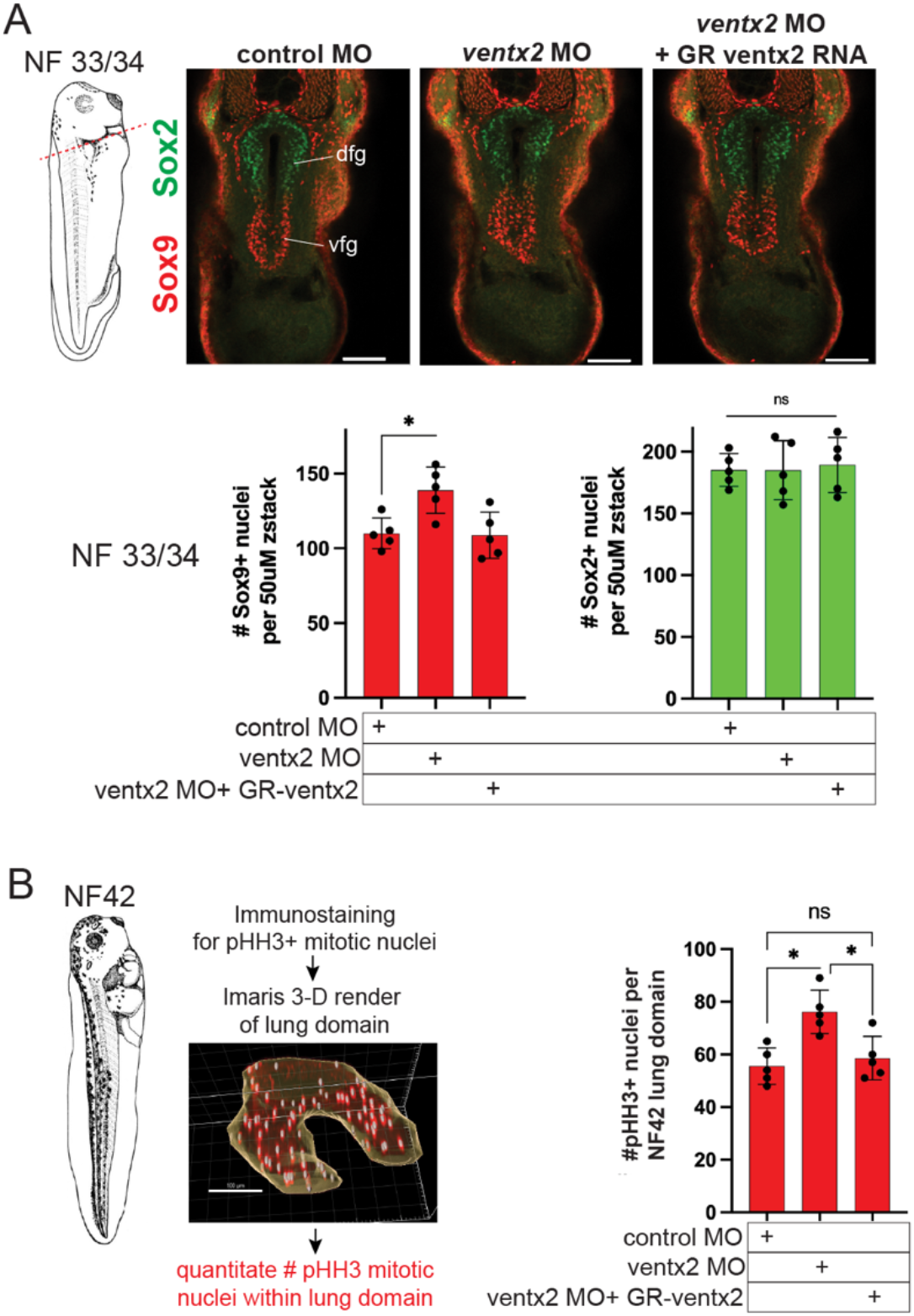
Quantitation of immunostaining for respiratory and esophageal progenitor number at NF33/34 and for mitotic phospho-Histone H3+ number within the NF42 lung domain. (A). Immunostaining panel is same as main Figure 4D. Quantitation was performed on these embryos, immunostained for the ventral respiratory progenitor marker Sox9 (red) and dorsal esophageal progenitor marker Sox2 (red). Numbers of positive nuclei per 50uM confocal z-stack are shown in the graphs; each black dot is a zstack from a separate embryo (n=5 per condition). Scale bar= 100uM. *Asterisk=p<0.05, parametric two-tailed T-test relative; ns=not significant. (B). Quantitation of phospho-Histone H3 (pHH3) positive nuclei within NF42 lung domain. Imaris software was used on embryos from figure 4C to generate 3-D renderings of the lung domain (example is shown) and within that domain the number of pHH3+ nuclei were quantitated. Each black dot is a separate embryo (n=5 per condition). Scale bar= 100uM. *Asterisk=p<0.05, parametric two-tailed T-test relative; ns=not significant.

**Supplemental Table S1.**
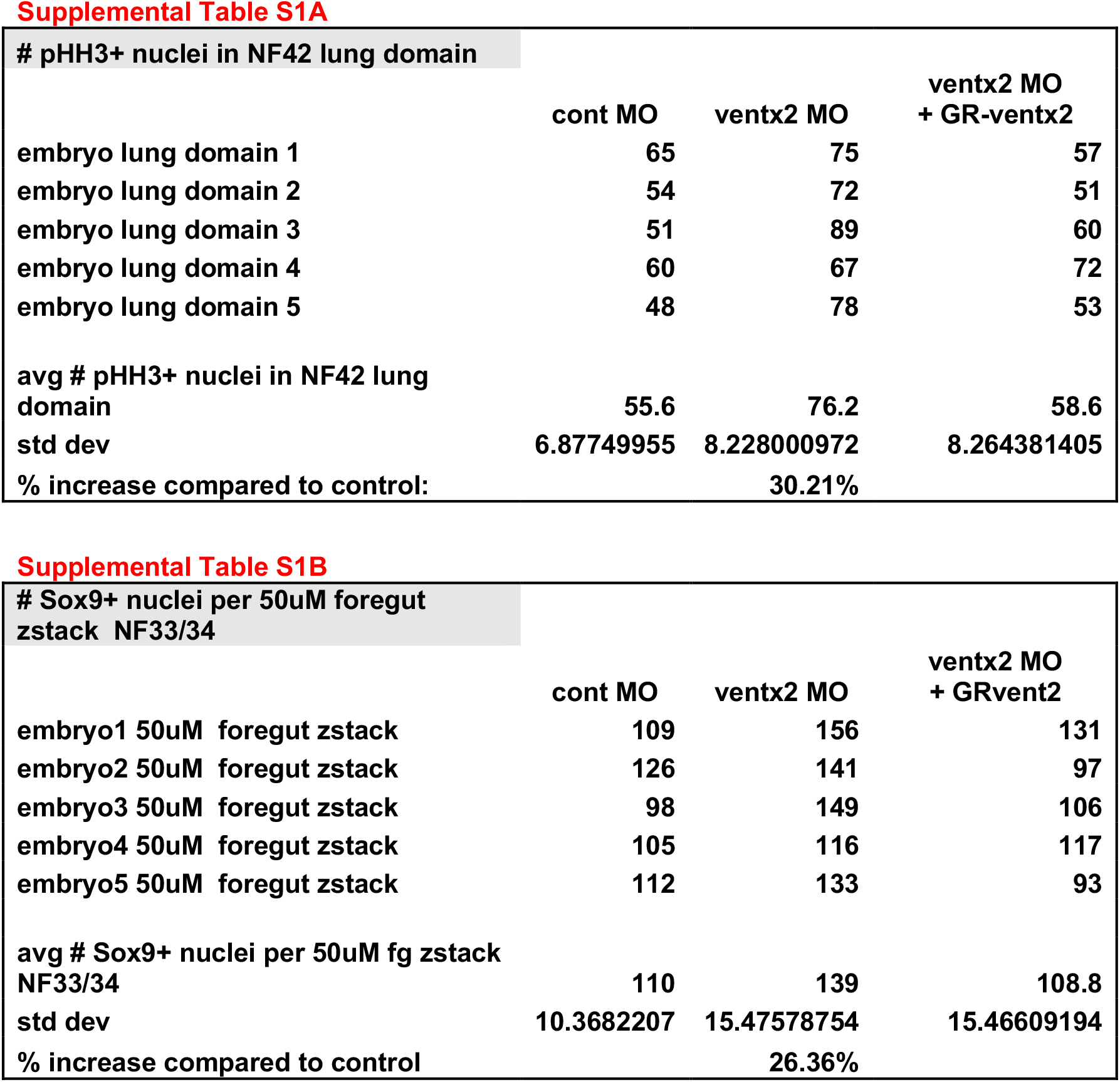
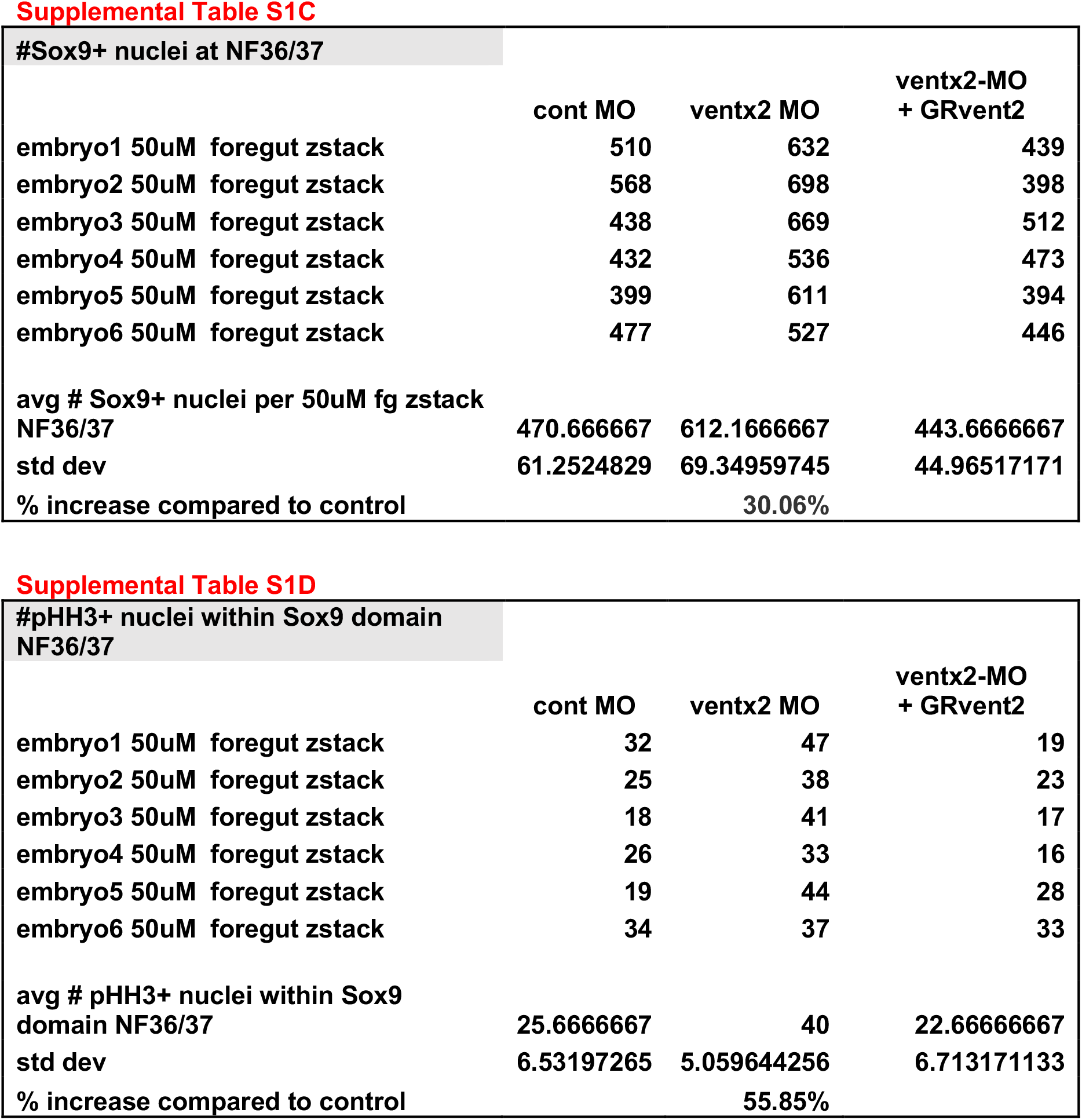
Quantitation of immunostaining data related to Figure 4 and Supplemental Figure S3.

## Notes

### Competing Interest Statement

The authors have declared no competing interest.

